# Digging roots is easier with AI

**DOI:** 10.1101/2020.12.01.397034

**Authors:** Eusun Han, Abraham George Smith, Roman Kemper, Rosemary White, John Kirkegaard, Kristian Thorup-Kristensen, Miriam Athmann

**Affiliations:** Department of Plant and Environmental Sciences, University of Copenhagen, Denmark; Department of Computer Science, University of Copenhagen, Denmark; Department of Agroecology and Organic Farming, University of Bonn, Germany; CSIRO Agriculture and Food, Canberra, Australia

**Keywords:** Deep Learning, Segmentation, Root quantification, Profile wall, Root washing, Soil coring

## Abstract

The scale of root quantification in research is often limited by the time required for sampling, measurement and processing samples. Recent developments in Convolutional Neural Networks (CNN) have made faster and more accurate plant image analysis possible which may significantly reduce the time required for root measurement, but challenges remain in making these methods accessible to researchers without an in-depth knowledge of Machine Learning. We analyzed root images acquired from three destructive root samplings using the RootPainter CNN-software that features an interface for corrective annotation for easier use. Root scans with and without non-root debris were used to test if training a model, i.e., learning from labeled examples, can effectively exclude the debris by comparing the end-results with measurements from clean images. Root images acquired from soil profile walls and the cross-section of soil cores were also used for training and the derived measurements were compared with manual measurements. After 200 minutes of training on each dataset, significant relationships between manual measurements and RootPainter-derived data were noted for monolith (R^2^=0.99), profile wall (R^2^=0.76) and core-break (R^2^=0.57). The rooting density derived from images with debris was not significantly different from that derived from clean images after processing with RootPainter. Rooting density was also successfully calculated from both profile wall and soil core images, and in each case the gradient of root density with depth was not significantly different from manual counts. Our results demonstrate that the proposed approach using CNN can lead to substantial reductions in root sample processing workloads, increasing the potential scale of future root investigations.

## Introduction

Destructive root measurements such as root extraction from soil samples, root counting from soil profile walls and cross-sections of soil cores are frequently used root methods *in situ* (Böhm 1976; van Noordwijk et al. 2001). Extracting roots from soil samples is commonly done for measuring root density and morphology. The extracted samples are scanned, and the acquired images are analyzed using generic software such as WinRHIZO (Regent Instruments Inc.). However, this procedure requires clean root samples without debris, this requires many hours of processing leading to downsizing the possible scale of the investigation.

Relatively faster measurement is possible by root-counting on soil profile walls. Experiments can involve 3-4 repeated measurements per season and may extend to soil layers below 1 m depth (Perkons et al. 2014; Han et al. 2015; Kemper et al. 2020). However, the ambiguity in root quantification using visual judgement with Root-Length Unit (1 RLU=0.5 cm) remains as a methodological challenge to generate robust end-results, and since these measures are estimates rather than precise root counts, results between different observers may vary in absolute terms.

The core-break method is a suitable and faster way to observe roots including their relation to soil structural features such as pores and cracks (White and Kirkegaard 2010). However, in common with the profile wall method, the root counts per unit core surface area is an estimated measure and extracting precise root length requires the time-consuming image annotation, or correlation with washed root samples as described above.

State-of-the-art Machine Learning approaches train Convolutional Neural Networks (CNN) based on user-machine interaction to generate models for fast and accurate segmentation - a set of pixels over the predicted target objects in real time (Kontogianni et al. 2020). CNN-based systems have demonstrated their potential to streamline the extraction of phenotypic traits from images for a broad range of tasks in plant science (Jiang and Li 2020) and have been found to be effective for segmentation of roots in both 3D X-ray CT images (Soltaninejad et al. 2020) and ordinary 2D photographs (Smith et al. 2020b).

Recently, using rhizotron, soil biopore and root nodule images, corrective annotation was found to facilitate accurate model training with annotations created within two hours and also for users without a background in Machine Learning (Smith et al. 2020a). This approach can replace time-consuming root extraction or counting with fully-automatic segmentation, leading to faster and more accurate results by root researchers. However, there have been few attempts to autonomously quantify roots using destructive sampling procedures (e.g. Teramoto and Uga 2020) as they may not include routine image acquisition or may comprise relatively few images in the dataset.

Therefore, we aimed to validate the use of an AI-based software called RootPainter (Smith et al. 2020a) for autonomous detection of roots using a corrective annotation strategy on images from incompletely extracted root samples, profile walls and cross-sections of soil cores.

## Materials and Methods

### RootPainter software

RootPainter is an image analysis software that enables rapid training of Deep Learning models with corrective annotations using a GPU server. RootPainter uses a modified U-Net (Ronneberger et al. 2015) architecture which is an encoder-decoder style Convolutional Neural Network (CNN) for semantic segmentation. It is a user-friendly software that features an interface for corrective annotation for efficient training even by non-specialists. A detailed description of the software is available in Smith et al. (2020a).

### Data acquisition

#### Monolith method

Soil monolith samples (10 x 10 x 10 cm in volume) from winter wheat (*Triticum aestivum* L.) were collected at Campus Klein-Altendorf (CKA) near Meckenheim, Germany (50°37’15’’ N 6°59’50” E), on a Haplic Luvisol in April 2020 at 10-20 cm, 40-50 cm and 60-70 cm soil depth. Root extraction was carried out by placing the sample on a sieve tower with sieve sizes 4 mm, 2 mm, 1 mm, 0.71 mm, 0.63 mm and 0.5 mm, and removing the soil with water. Afterwards, roots and remaining debris of individual sieves were photo-scanned using Epson V700 at a DPI of 800 in TIFF format. The original scans were converted to JPEG at the same DPI for training. After scanning, the root samples were further cleaned by removing remaining debris, and then scanned using the same procedure.

#### Profile wall method

Soil profile wall images from a field trial with different fodder crops, namely, lucerne (*Medicago sativa* L.), tall fescue (*Festuca arundinacea* Schreb.) and a mixture of lucerne and tall fescue were acquired at Hofgut Oberfeld near Darmstadt, Germany (49°52’53”N 8°42’02”E), on a Gleyic Luvisol, in September 2020. The fodder crops had been sown in spring 2019. Images were taken from a profile wall flattened with spades, from which 0.5 cm of soil were carefully removed using a fork, from 0-100 cm of soil depth. Each image was 50 cm x 50 cm in size. The raw images were acquired using Nikon 7100D with Nikon AF-Nikkor (16-85 mm) lens having 18 mm focal length and 13 aperture. The distance between the profile walls and the lens was 80 cm. The camera produced NEF format images which were further converted to JPEG with 1200 DPI. From the same profile walls, manual counts were recorded in terms of Root-Length Unit (RLU) equivalent to 0.5 cm length.

#### Core-break technique

Soil core samples were taken from a standing wheat crop at anthesis on a red Kandosol soil near Bethungra, NSW, Australia in 2005 (34°43’S, 147°48′E). The detailed description of the site and the acquisition and processing of the images is available in White &Kirkegaard (2010). Briefly, the soil cores were collected at depths of 0-40 cm, 40-80 cm, 80-120 cm and 120-160 cm using thin-walled steel tubes 9.4 cm in diameter and 40 cm in length. The collected cores were broken transversely at 7-15 cm and 4-12 cm from the upper and lower surfaces. Using A Zeiss Axiocam digital camera with Axiovision software (Carl Zeiss Australia, Sydney, Australia), 10 random images of 1 cm2 area from the revealed surface areas were captured at high resolution (2600 x 2060 pixels). For the images not in focus, using AutoMontage Essentials (Syncroscopy, Frederick, MD, USA), they were combined into a single image with most regions of the image in focus. The raw data on root counts were provided by the authors.

### Training and analysis

Training datasets consisting of 648 (975 x 1000 pixels), 4240 (700 x 700 pixels) and 239 (1200 x 951 pixels) images were prepared from root scans after extraction from the monoliths, soil profile walls, and cross-sections of soil cores, respectively. Annotation/training time for each dataset was restricted to 200 minutes. Instructions for server setup, software installation and corrective annotation are available from Smith et al. (2020a). At least 6 images showing clear objects (roots) of interests were annotated without the prediction from CNN, after which corrective annotation – annotating based on comparing CNN model prediction with annotator observation on the image was initiated. After completion of each training, the final models were used to generate segmentations on the original images in PNG format. The segmented images were used to compute the length of roots as a sum of the pixels of the predicted roots after skeletonization as described in Smith et al.

#### Monolith method

After training, the segmented images from both completely and partially extracted root samples from the monoliths were fed to ‘WinRHIZO pro’ (Version 2019, 32bit) in JPEG format, from which root length (cm) was generated. Root length was converted to root-length density (RLD; cm cm^-3^) by dividing the root length by the soil volume (10 x 10 x 10 cm). RLDs derived from segmentation on partially and fully cleaned samples and from WinRHIZO analysis on fully cleaned samples were used for linear regression analysis. Additionally, segmented images on the fully cleaned samples which were fed to WinRHIZO analysis were also included for mean comparisons of RLDs (Tukey HSD, P≤0.05).

#### Profile wall method

The segmented images from profile walls were split into 10 x 10 sub-images to match the original regions for manual counts (5 x 5 cm), on which linear regression was performed. RLDs from segmented images and manual root counts were calculated from RLU and assumed soil volume (5 x 5 x 0.5 cm), which were used for mean comparisons (Tukey HSD, P≤0.05) and linear regression analysis.

#### Core-break technique

For the segmented images from the core-break method, root counts from the original study (White and Kirkegaard 2010) were used for validation. However, unlike the analyses done with root extraction and profile wall counting, there was no data on root length, which did not allow direct comparisons of root length between segmentation and manual measurements. Therefore, we focused on comparing their differences along the soil depth by performing mixed-effects model analysis (P≤0.05) followed by multiple comparisons (Tukey HSD, P≤0.05).

Statistical analyses were done using R (R Core Team 2019). For linear regression **lm** function was used and **lmer** function was used for mixed-effects model analysis.

## Results and Discussion

### Training performance

After providing the first 6 examples, CNN predicted the roots fairly accurately from the background from the scans of partially cleaned roots, however, the debris were also segmented as roots. From the 7^th^ images after the corrective annotation (13^th^ image in total) the software began to exclude the debris. Further refining on the root boundaries with the remaining images up to the 355^th^ image within 200 minuites yielded a satisfactory model not only visually (Figure 1a) but also by linear regression between manual measurements by WinRHIZO on cleaned samples and segmentation on samples with debris (R^2^=0.99; Figure 2a).

**Figure 1.**
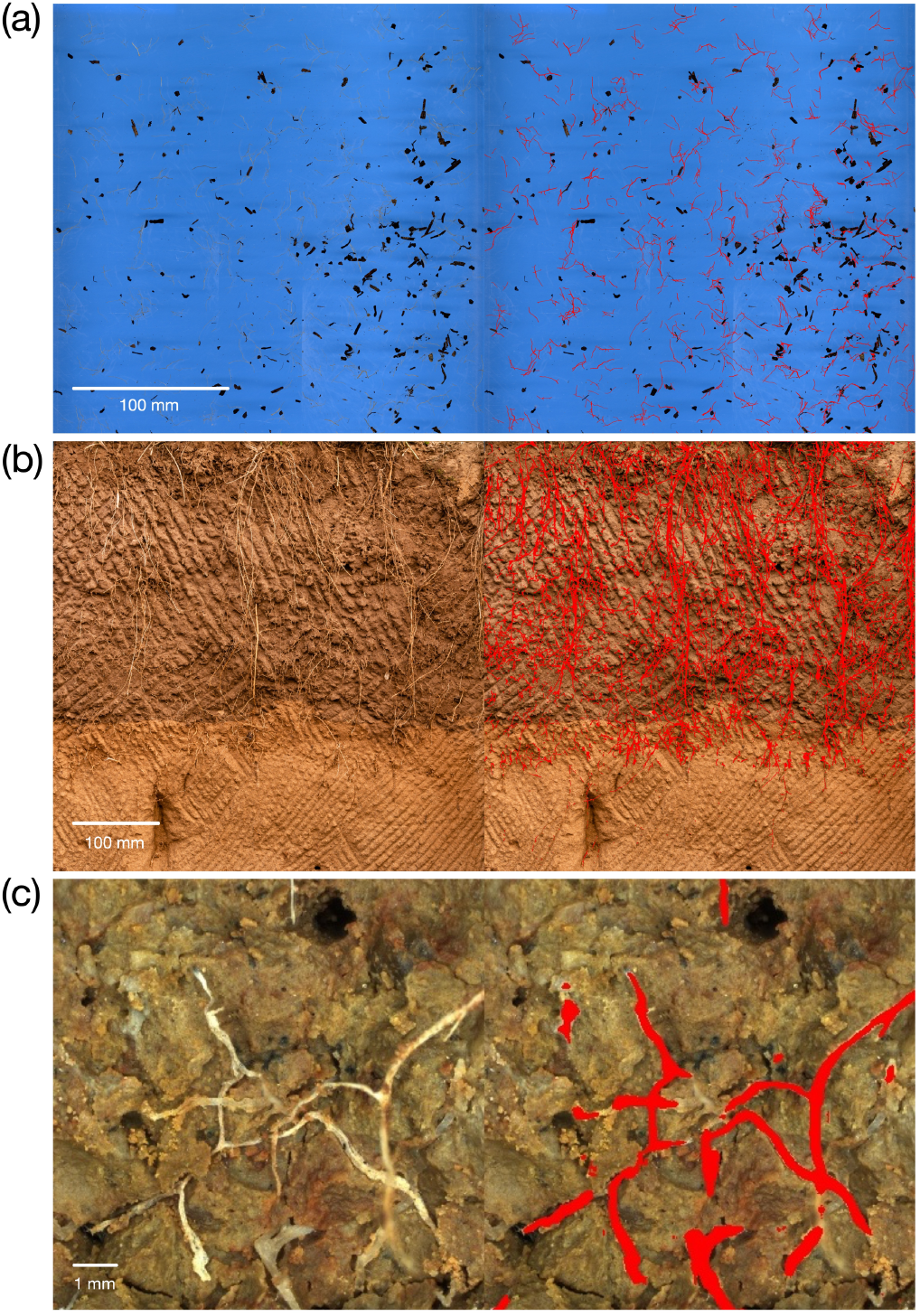
Comparative view on original and trained images of roots from soil monolith (a), profile wall (b) and core-break section (c). (2020b).

**Figure 2.**
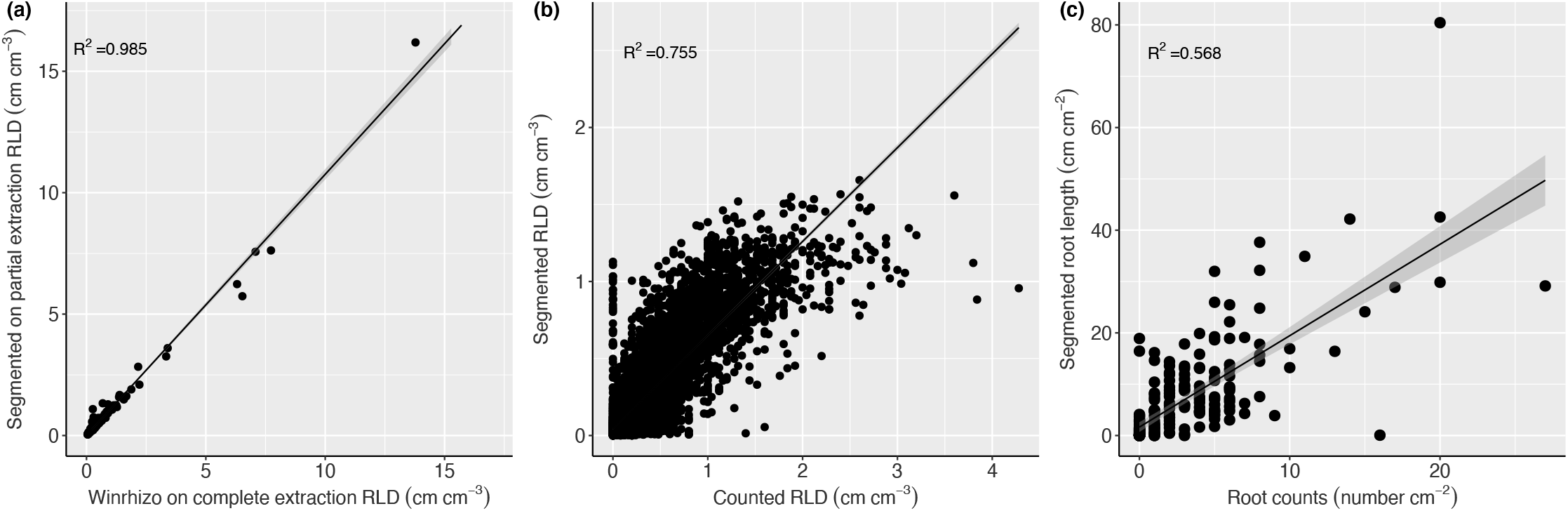
Linear regressions between manual measurements and AI segmentation on soil monolith (a), profile wall (b) and core-break methods (c).

The high accuracy on extracted root scans despite the debris may be attributed to the relatively uniform background resulting from the consistent scanning quality with high DPI (800 DPI). Preparing high quality images is one of the prerequisites for obtaining accurate results from Deep Learning models (Dodge and Karam 2016), and root scans from the Epson V700 can result in optimal images for machine training. However, such images, especially uncompressed RGB images can increase the demand for storage, transfer time and can be more cumbersome to work with in the user-interface. Using RootPainter we were able to use the in-built functionality to create a training dataset consisting of images with smaller pixel numbers (950 x 1000 pixels), which enhanced the speed of the image-loading, but segmentation and data extraction using the original images required more time to complete compared to the other types of images used in this study.

The relationship between the RLDs derived from manual counts and from the segmentation on soil profile walls was significant with an R^2^ of 0.76 (Figure 2b). Training on the profile wall images was more challenging due to the varying soil background. For example, pore channels and disturbed soil surfaces made it difficult to accurately segment the visible roots at the early stage in the interactive training procedure (Figure 1b). Moreover, the original images contained 3300 x 3300 pixels covering the area of 50 x 50 cm (≈44 pixels mm^-2^). This is approximately 23 times lower in resolution compared with the images from the root scans (≈996 pixels mm^-2^) which led to blurring of target roots, which may have affected the model performance (Dodge and Karam 2016). Moreover, on the topsoil region, the hanging aboveground shoots caused false positives, which should be avoided during image acquisition or be removed carefully during dataset preparation. However, we believe that this is still a promising result as, firstly, the counter and image annotator were two different individuals. Secondly, the high R^2^ from 5 x 5 cm-wise linear regression implies that the segmented images can be used for repeated analysis on the soil depth and horizontal replicates which provides robust results from statistical analysis as shown previously (Han et al. 2015; Kemper et al. 2020).

Linear regression analysis between the original root counts by the core break method and segmentation on the stitched images from soil cores revealed the R^2^ of 0.57 (Figure 2c). Interactively training a model to segment the cross-section images from soil cores was particularly challenging, judging by the high degree of false positives observed during the training procedure. This may be due to fewer “roots” in the images, which covered an area of only 1 cm^2^, and that some samples were taken from deep soil layers (up to 160 cm of soil depth), which generally contain few roots. A further bottle-neck for improvement of predicted root profiles may have arisen from the image stitching used to combine extended focus images, as several images showed local distortions (Figure 1c). Finally, the lower performance may have stemmed from performing regression analysis between distinctively different variables with the core break method, i.e, root counts and root length.

### Manual measurements vs. segmentation

Mean comparison between RLDs analyzed by WinRHIZO on clean images, segmented images without debris and segmented images with debris revealed no significant differences at the measured soil depths (Figure 3a). This is the first result demonstrating the compatibility of AI-segmentation to streamline further analysis of root parameters, e.g. through WinRHIZO analysis, which can allow further analysis for root morphology with detailed root diameter classes (Han et al. 2016). However, based on our preliminary analysis, the segmentation led to an underestimation of the finer roots smaller than 0.1 mm compared with generic WinRHIZO analysis with clean samples (data not shown). More detailed size classification should be used to determine the degree of morphological shift from generic analysis to AI-segmentation for confirmation. Alternatively, analyzing the segmentation using open source software with publicly available source code such as RhizoVision Explorer (Seethepalli et al. 2020) would allow a more thorough investigation into the cause of discrepancies in fine root quantification.

**Figure 3.**
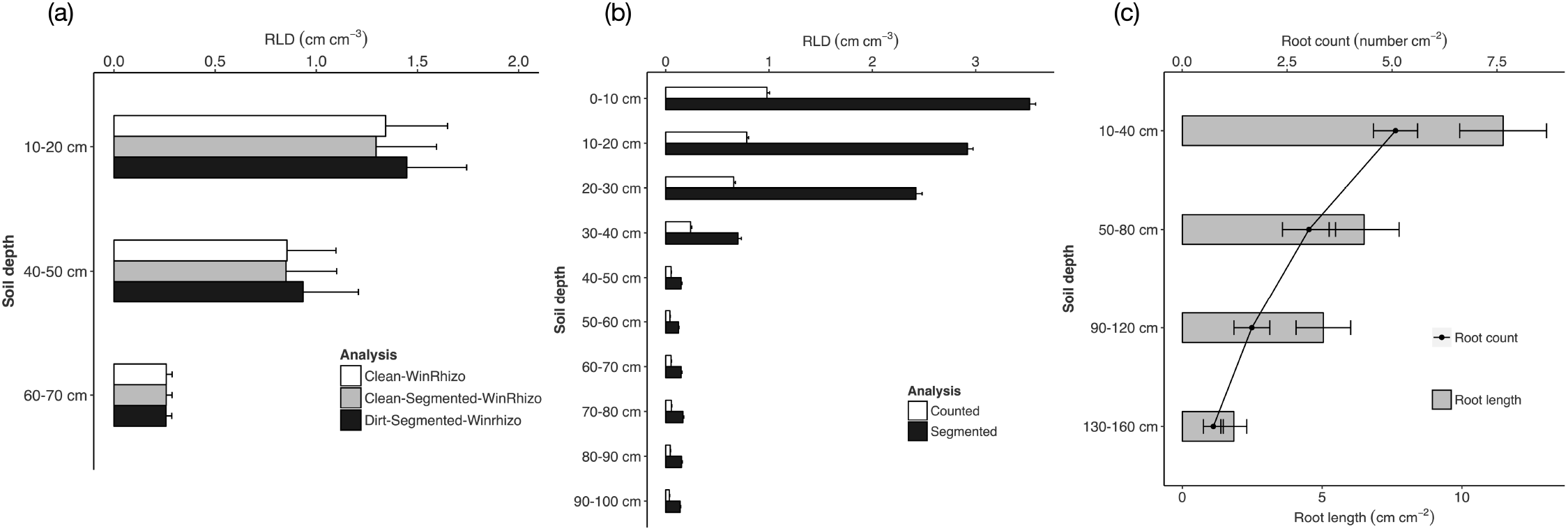
Mean comparisons between generic measurements and AI segmentation on soil monolith (a), profile wall (b) and core-break methods (c).

We believe that our approach is suitable especially for those measuring root density with a large number of soil samples such as for breeding research programs (Wasson et al. 2014). According to our experience, manual removal of debris takes about 3 or 15 standard working hours for subsoil and topsoil samples of 1000 cm^3^, respectively. If it can be omitted, in our estimate only 50 % of the time for subsoil samples and only 10 % of the time for topsoil samples will be required for processing.

Mean comparison between manual counts and segmentation from the profile wall method resulted in significant differences in RLD at all the soil depths, except for 90-100 cm (Figure 3b). Consequently, we are cautious regarding the accuracy of the absolute RLD. However, given that the root counters used an abstract scale of 0.5 cm to measure RLU, whereas the RLD from the segmentation was derived from pixel-scale, we assume that the latter represents more accurate root measurement. Visual judgement used for the profile wall method is a bottle-neck and can underestimate the rooting density, especially in the topsoil layers where roots are densely growing. When compared between the soil depths, the two RLDs derived from manual counts and segmentation resulted in the same effects – the highest RLD was observed at 0-10 cm and decreased with depth. While the significant differences in predicted RLD were shown the top 5 soil layers, below 40-50 cm there were no significant differences evident. Further validation using soil sampling as a calibration factor can lead to a firm conclusion, which is a common practice for the profile wall method (Perkons et al. 2014).

The profile wall method is commonly used to investigate root growth dynamics with multiple observation points during the season (Kautz et al. 2013). However, the number of replicates investigated is often restricted, for example, to two plots per treatments due to time constraints (Han et al. 2017). By adopting automatic segmentation, the counting time (1-4 hours per plot), which was the main bottle-neck of the profile wall observation, can be eliminated, allowing more treatments or larger areas to be monitored (e.g. transects in agroforestry systems).

Due to the absence of measured root length, we could not directly compare the root density in the same units between the segmentation to the original data from White & Kirkegaard (2010) on the core-break method. Nevertheless, the patterns of root length at depth derived from segmentation was identical to the original root counts reported in White & Kirkegaard (2010). For both measures the highest rooting density was noticed at 10-40 cm followed by 50-80, 90-120 and 130-160 cm. Therefore, we conclude that the model trained with RootPainter provided biologically meaningful results despite the weak correlation with manual counts. Soil core sampling often involves multiple purposes such as water/nutrient measurement (Lilley and Kirkegaard 2011) and biopore/crack quantification (White and Kirkegaard 2010). With a more standardized image acquisition process, autonomous segmentation could lead to faster and more accurate root measurements, which can be easily related with other parameters acquired from the same samples.

Our results suggest that the time for root processing from soil samples, and also for root counting on soil profile walls and on soil cores can be substantially reduced and more accurate end-results can be obtained using human-machine interactive training. Therefore, we conclude that automatic segmentation with Deep Learning can contribute to efficient root quantification in field conditions using destructive procedures leading to reduced time and cost for extracting and measuring. This will lead to lower demand for time and cost for counting, annotating and extracting root samples, which can allow us to extend the scale of research objectives in field-based root studies.

## Availability of data and materials

The dataset and manual counts used in the study are available online (Han et al. 2020b). The created training dataset and final trained models are available online (Han et al. 2020a). The python script for splitting the segmentation on profile wall images is available from Smith (2020).

## Author contributions

All authors contributed to manuscript writing. EH and MA co-designed the research. EH conducted the research and prepared the manuscript. AGS provided technical advice on Machine Learning and wrote a python script for splitting the segmented images. RK organized the data on manual counts from German sites. RW and JK provided a guidance for recognition of the Australian biopores and performed a quality check on segmentation. The GPU server was provided by KTK. MA coordinated the root sampling and data collection at German sites.

## Acknowledgments

The soil monoliths were taken and processed within the BonaRes research project Soil^3^ (grant number 031B0515E) funded by the German Federal Ministry of Education and Research (BMBF). The profile wall images were generated within the project, MIKODU’ (grant number 2818OE024) supported by funds of the German Federal Ministry of Food and Agriculture (BMEL) based on a decision of the parliament of the Federal Republic of Germany via the Federal Office for Agriculture and Food (BLE) under the Federal Programme for Ecological Farming and Other Forms of Sustainable Agriculture. Use of GPU server for Machine Learning was supported by the project DeepFrontier (funded by Villum Foundation; grant number VKR023338). Special thanks go to Johannes Siebigteroth and Christian Dahn for preparing profile wall images for annotation, and to Ariane Eckstein and Stefanie Fuchs for their extra work with scanning the soil monolith samples.

